# Altered placental morphology and metabolomic profile in association with uncomplicated metabolically healthy obese pregnancy

**DOI:** 10.1101/2025.03.20.644371

**Authors:** Ousseynou Sarr, Akasham Rajagopaul, Shuang Zhao, Xiaohang Wang, David Grynspan, Genevieve Eastabrook, Liang Li, Timothy R.H. Regnault, Barbra de Vrijer

**Author notes:** These authors contributed equally to this work and share senior authorship (Timothy R.H. Regnault, Barbra de Vrijer).

## Abstract

Metabolically healthy obesity (MHO) and metabolically unhealthy obesity (MUO) in pregnancy represent two distinct cardiometabolic populations, each potentially necessitating alternative clinical management. However, the understanding of the unique physiological effects of uncomplicated MHO or MUO on fetoplacental growth and metabolism remains limited. This study aimed to identify changes in placental morphology and metabolites concerning maternal obesity, independent of pregnancy-related cardiometabolic complications. Placentae from women with a pre-pregnancy body mass index (BMI) <25 kg/m^2^ (control; n = 15) and women with MHO (pre-pregnancy BMI >30 kg/m^2^ with no cardiometabolic diseases; n = 15) were analysed for indices of placental growth and untargeted metabolomics. Complementary measures of pro-inflammatory and antioxidant genes, as well as antioxidant proteins, enzymes, and lipid peroxidation markers were conducted. A clear placentomegaly without histopathological changes was observed in uncomplicated MHO pregnancies. The metabolite 3-aminoisobutanoic acid emerged as the top-ranked feature distinguishing MHO from control placentae, and cysteine, methionine, and vitamin B6 metabolism pathways were among the most distinct signatures identified. These findings illustrate an altered placental morphology and metabolomic profile specific to uncomplicated MHO, offering new insights into how obesity without cardiometabolic complications influences fetoplacental growth and metabolism. They also represent a crucial first step towards marker identification for MHO pregnancies and underscore the importance of alternative care pathways when obesity is present but metabolic comorbidities are absent.

## 1. Introduction

Obesity, defined as an abnormal or excessive body fat and measured by a body mass index (BMI) ≥ 30 kg/m^2^, is traditionally associated with adverse metabolic health outcomes and has become a global epidemic, posing a significant challenge to public health [1,2]. It is linked to a variety of metabolic and cardiovascular complications, including insulin resistance, type 2 diabetes, high blood pressure, and abnormal blood lipid levels [1]. However, evidence indicates that obesity does not always lead to adverse metabolic outcomes since BMI *per se* does not take into account the heterogeneity of body fat distribution, which is a key driver of cardiometabolic risk associated with obesity [3,4]. Around 10-30% of people with obesity possess a metabolically healthy status, that is, the absence of overt cardiometabolic abnormalities [2]. This concept is referred to as ‘metabolically healthy obesity’ (MHO), in contrast to a metabolically unhealthy state, characterized by the presence of complications such as hypertension or diabetes (metabolically unhealthy obesity; MUO) [2]. Although there is no standardized definition of MHO, it generally refers to the absence of metabolic and cardiovascular diseases, including type 2 diabetes, dyslipidemia, hypertension, insulin resistance and atherosclerotic cardiovascular disease in a person with obesity (BMI ≥30 kg/m2) [2,5].

Obesity complicating pregnancy particularly poses a concern, as an escalating number of women of reproductive age are now being categorized as obese [6]. It is estimated that obesity affects up to 25% of pregnant women in the Western World [7]. Maternal obesity is linked to a higher likelihood of experiencing obstetrical complications, including gestational diabetes mellitus (GDM), gestational hypertension, preeclampsia, preterm delivery, large for gestational age (LGA) baby, caesarean section, as well as an increased risk of neonatal morbidity and mortality [8]. In addition to potentially experiencing negative immediate consequences, both the mother who has obesity and her child are at an increased risk of developing cardiovascular, metabolic, and neurological disorders post pregnancy in later life [9–12]. During pregnancy, maternal obesity with a metabolic comorbidity is referred to as MUO, similar to the non-pregnant population [9]. However, it is important to note that not all pregnant women with obesity experience metabolic dysfunction. In fact, a significant number are classified as having MHO [13] and do not exhibit cardiometabolic complications typically associated with maternal obesity [13–15], even though they deliver LGA babies [16] and an increased placental size, similar to MUO pregnancies [16,17].

The placenta plays a crucial role in the development and growth of the fetus, and changes in placental development and growth are associated with poor maternal, newborn and offspring outcomes [18,19]. Indeed, changes in measures of placental growth including birth weight/placental weight ratio (BPW) and birth weight/placental volume ratio (BPV), measures of placental insufficiency, are observed in conjunction with adverse outcomes such as preeclampsia and GDM, and can be used as predictors of pregnancy complications [20–22]. These associations are different from those of fetal birth weight or placental weight alone [22]. Moreover, these measures of the placenta itself have been associated with altered metabolism and as indicators /readouts of the in-utero environment. Placental breadth is associated with neonatal body size and may be an important indicator of placental nutrient transport capacity [23]. Additionally, the surface area and thickness of the placenta are proportional to the number of uterine spiral arteries that provide placental oxygenation and nutrition, and the extent of villous branching, respectively [24–26]. Furthermore, in the pregnancy with both obesity and GDM, the placenta is relatively large for fetal size, resulting in a relatively small birth–placental weight (BPW) ratio, suggestive of a reduced placental efficiency in pregnancies with obesity [27]. These placental morphometric changes, reflective of the in-utero environment, are also likely related to changes in placental nutrient transporter expression, mitochondrial function, lipid metabolism, and oxidative stress levels [28]. This change in the birth-placental weight ratio may be influenced by placental inflammation, oxidative stress, and shifts in placental metabolism linked to obesity itself—specifically metabolically healthy obesity (MHO)— regardless of the cardiometabolic comorbidities observed in metabolically unhealthy obesity.

The metabolome, the complete set of small-molecule chemicals (metabolites) found within a cell, cellular organelle, an organ, tissue, or biofluid, is best investigated through a comprehensive approach named metabolomics [29]. Metabolomics is used to identify phenotypical groups with specific metabolic profiles, and its application to various pregnancy conditions is on the rise [30]. Studies of placental metabolomics during uncomplicated and complicated obese pregnancies with optimal prenatal care (specific nutritional advice and recommendations on weight gain in pregnancy), have identified specific metabolomic signatures of the placentae from people with obesity [31,32]. These studies mostly investigate clinical obesity in general, without differentiating between metabolically healthy and unhealthy phenotypes. A better understanding of how obesity without comorbidities (i.e., uncomplicated MHO) may alter placental morphology and the metabolomic profile could help define baseline changes in placental growth and function, as knowledge in this area is still evolving [13,15,16].

This study accordingly examines the changed placental morphology and metabolomic profile related to uncomplicated MHO pregnancy. The primary aim was to analyze placental growth and metabolites at term in uncomplicated MHO pregnancies using liquid chromatography–mass spectrometry (LC–MS) and to evaluate placental changes in antioxidant, pro-inflammatory genes, and lipid peroxidation markers in comparison to healthy controls. It was hypothesized that these findings would clarify how obesity without cardiometabolic comorbidities affects fetoplacental growth and metabolism. Furthermore, they could provide an essential initial step in identifying metabolomic markers for MHO and in guiding alternative clinical care pathways for pregnancies complicated by obesity in the absence of overt metabolic dysfunction.

## 2. Materials and Methods

### 2.1. Population

The protocol of the study was approved by the Western University’s Human Subjects Research Ethics Board (REB# 106663). Pregnant women (normal weight; 18≤ pre-pregnancy BMI <25 kg/m^2^, (control (C); n=15) and with obesity; pre-pregnancy BMI >30 kg/m^2^ (MHO), n=15) were recruited at their planned caesarean delivery without labor at term (>37 weeks’ gestational age, GA). Exclusion criteria included multiple pregnancy, age <18 or >40 years at estimated due date, maternal hypertensive disorders of pregnancy, gestational diabetes or other significant medical conditions predisposing to placental dysfunction (renal, autoimmune disease), pre-pregnancy BMI <18 kg/m^2^ or pre-pregnancy BMI between 25 and 30 kg/m^2^. Hypertension in pregnancy was defined as systolic blood pressure (BP) of >140 mmHg and/or diastolic BP of >90 mmHg at least 2 instances after 20 weeks of pregnancy in pregnancies with normal initial blood pressures. Preeclampsia was defined as de novo hypertension after 20 weeks of gestation in combination with proteinuria. Gestational diabetes was excluded with a glucose challenge or tolerance test (GCT/GTT) during the second trimester. Maternal clinical characteristics, obstetrical and neonatal outcome data were collected and have been previously described by Cohen et al. [16]. As per the exclusion criteria and maternal clinical characteristics, the group with obesity was classified as having metabolically healthy obesity (MHO), representing pregnancies without the typical complications associated with obesity, such as hypertension, gestational diabetes, or other related conditions.

### 2.2. Placenta tissue collection

Placentae were collected at elective caesarean sections and placental weight, length, thickness, breadth and surface area were recorded. Placental tissue samples (n=15 in each group) were collected from three different areas [centre, periphery (1–2 cm from the placental edge), and middle (between these two sections)] within 30 min of delivery, snapfrozen in liquid nitrogen and stored at −80°C as previously reported [16].

### 2.3. Placental histology

Formalin-fixed paraffin-embedded placental tissues were sectioned and stained with hematoxylin and eosin (Molecular Pathology, Robarts Research Institute). Stained slides were scanned using Aperio ScanScope CS (Leica, Microsystems CMS GmbH, Wetzlar, Germany). Slides of the peripheral, middle, and central regions of each MHO and control placenta were examined by a clinical pathologist (DG), in a blinded manner, and histopathological features recorded and graded according to the following scoring system: distal villous hypoplasia (none, focal or diffuse), increased syncytial knots (none or present), chorangiosis (none or present), delayed villous maturation (none, focal or diffuse), avascular fibrotic villi (none or small, intermediate or large foci), increased focal perivillous fibrin deposition (none or present), massive perivillous fibrin deposition (none or diffuse), maternal floor infarct pattern (none or present), intervillous thrombi (none or present), and chronic inflammation (none, low-grade or high-grade). The distributions of these different placental histopathological features are represented in frequency and percentage as described previously [33].

### 2.4. Untargeted metabolomics using chemical isotope labeling (CIL) GC-MS

All chemicals and reagents were purchased from Sigma-Aldrich (Markham, ON, Canada), except those specifically stated. Liquid chromatography-mass spectrometry (LC-MS) grade water, acetonitrile, methanol and formic acid were purchased from Canadian Life Sciences (Peterborough, ON, Canada). For in-depth placenta metabolomics, a high-performance LC–MS was performed at The Metabolomic Innovation Centre (TMIC), the Department of Chemistry, University of Alberta (Edmonton, AB, Canada) according to the published analytical workflow [34]. Placental samples from the different placental regions were individually processed during the analysis.

Placental samples were homogenized in methanol, dichloromethane (DCM) and water and metabolites extracted as previously described [35]. Samples were randomized before any procedures to eliminate any potential batch variations in sample analysis. Sample normalization was carried out by measuring the total metabolite concentration in each sample[36,37]. A proprietary metabolome quantification kit from Nova Medical Testing Inc. (Product Number: NMT-6001-KT) was used to measure the total concentration in an aliquot of 25 µL of sample extract. Water was added to adjust all the concentrations of samples to 1.2 mM.

Chemical isotope labeling of each sample was carried out by following the SOPs provided in the labeling kit (Nova Medical Testing Inc., Product Number: NMT-4101-KT. For labeling experiment, an aliquot of 25 µL of sample was used. Using this kit, the amine/phenol metabolites were labeled using dansylation reaction [38].

After labeling, the ^12^C_2_-labeled individual sample was mixed with ^13^C_2_-labeled reference sample in equal volume. The mixture was then injected into LC-MS for analysis. Prior to LC-MS analysis of the entire sample set, quality control (QC) sample was prepared by equal volume mix of a ^12^C-labeled and a ^13^C-labeled pooled sample. QC samples were run at an interval of one QC injection after 10 sample injections. All LC-MS analysis were carried out on using Agilent 1290 LC linked to Bruker Impact II QTOF Mass Spectrometer. The column used was Agilent eclipse plus reversed-phase C18 column (150 × 2.1 mm, 1.8 µm particle size) and the column oven temperature was 40 °C. Mobile phase A was 0.1% (v/v) formic acid in water and mobile phase B was 0.1% (v/v) formic acid in acetonitrile. The gradient setting was t = 0 min, 25% B; t = 10 min, 99% B; t = 15 min, 99% B; t = 15.1 min, 25% B; t = 18 min, 25% B. The flow rate was 400 µL/min. Mass spectral acquisition rate was 1 Hz, with an m/z range from 220 to 1000.

All raw LC-MS data were first converted to .csv files using DataAnalysis 4.4 (Bruker Daltonics, Bremen, Germany). The exported files were uploaded to IsoMS Pro 1.2.12 (Nova Medical Testing Inc.) for data processing and metabolite identification. ^12^C-/^13^C-peak pairs in each sample were first extracted and peak intensity ratio was calculated for each peak pair [39]. In this step, all redundant information was filtered out (e.g., adduct ions, dimers, etc.) to retain one peak pair for each metabolite. Then the same peak pair (i.e., metabolite) from different samples were aligned and the missing ratio values were filled back by the software. Data cleansing was carried out to remove peak pairs that originated from blank samples and that were not presented in at least 80.0% of samples in any group. Data was then normalized by Ratio of Total Useful Signal, calculated as sum of all useful ^12^C-peaks over sum of all useful ^13^C-peaks, served as post-acquisition normalization [37].

Metabolite identification was carried out using a three-tiered approach against NovaMT Metabolite Databases 2.0 (Nova Medical Testing Inc.) [35]. In tier 1, peak pairs were searched against a labeled metabolite library (CIL Library) based on accurate mass and retention time, considering as positive identification. In tier 2, the remaining peak pairs were searched against a linked identity library (LI Library), which includes over 9,000 pathway-related metabolites, providing high-confidence putative identification results based on accurate mass and predicted retention time matches. In tier 3, any remaining peak pairs were searched, based on accurate mass match, against the MyCompoundID (MCID) library (www.mycompoundid.org) composed of 8,021 known human endogenous metabolites (zero-reaction library), their predicted metabolic products from one metabolic reaction (375,809 compounds) (one-reaction library) and two metabolic reactions (10,583,901 compounds) (two-reaction library) [40].

### 2.5. RNA extraction and real time PCR

Placental tissue samples from the center, periphery, and middle of the placenta were pooled within each subject and ground prior to real-time PCR and spectrophotometric analyses. Total RNA was extracted from 50 mg of placental tissue using 1 mL of Grizol (Invitrogen, Burlington, ON, Canada) and the E.Z.N.A Total RNA kit I, following the manufacturer’s instructions (OMEGA bio-tek, Norcross, GA, USA). The quantity of extracted RNA was assessed using a Nanodrop spectrophotometer (Thermo Fisher Scientific, Waltham, MA), and RNA quality and integrity were evaluated via the A260/A280 ratio and agarose gel electrophoresis. One microgram of RNA was reverse transcribed into cDNA using the M-MLV Reverse Transcriptase Kit (Thermo Fisher Scientific) following the manufacturer’s instructions. Real-time PCR was performed using the SensiFAST SYBR No-ROX Kit (FroggaBio, Concord, ON, Canada), and all primers (**Supplementary Table S1**) were purchased from Sigma-Aldrich (Oakville, ON, Canada). The specificity of the PCR products was assessed using melting curve analysis. Non-transcribed RNA and PCR reactions using water instead of template showed no amplification. Gene expression analysis was conducted using the CFX Maestro software (Bio-Rad, Mississauga, ON, Canada) with the 2^-ΔΔCt method [41]. Beta-actin and proteasome 20S subunit beta type 6 (PSMB6) genes were used as reference genes. The average Ct values of the target gene were normalized to the average beta-actin and PSMB6 Ct values of the same cDNA sample. Beta-actin and PSMB6 were determined to be suitable reference genes based on the GeNorm algorithm in the CFX Maestro software, with an average expression stability (Avg M value) of 0.68 for both genes.

### 2.6. Antioxidant marker assays

Placental tissues (50 mg) were placed in ice-cold homogenization buffer (20 mM HEPES, 70 mM sucrose, 1 mM EGTA, and 210 mM mannitol). Each placental sample was homogenized in 500 μL of isolation buffer using a PRO Scientific Bio-Gen PRO200 Homogenizer. The resulting homogenate was divided into 2 parts and centrifugation at different speeds yielded the following supernatants: supernatant 1 containing superoxide dismutase (SOD) at 1,500g for five minutes at 4 °C; supernatant 2 containing catalase (CAT) at 10,000 g for 15 minutes at 4 °C. SOD (kit no. 706002, Cayman Chemical, Ann Arbor, MI, USA) and CAT (kit no. 707002, Cayman Chemical) enzyme activities, in placental homogenates were measured by spectrophotometric analysis following the kit instructions. Reduced glutathione (GSH), oxidized GSH (GSSG) and the GSH/GSSG ratio were determined in 50 mg of placental tissue homogenised in 1 mL of 2-(N-morpholino)ethanesulphonic acid (MES) buffer provided in the GSH assay (kit no. 703002, Cayman Chemical) and following the kit instructions.

### 2.7. Measurement of lipid peroxidation markers

Total 8-isoprostane levels in the placental tissues were measured using the 8-isoprostane enzyme-linked immunosorbent assay kit (kit no 516351, Cayman Chemical), following the manufacturer’s instructions. Briefly, 50 mg placenta samples were homogenized in 500 μL of 0.1 M phosphate buffer, pH 7.4, containing 1.4 mM EDTA and 0.005% butylated hydroxytoluene (BHT), using a PRO Scientific Bio-Gen PRO200 Homogenizer. After homogenization, samples were centrifuged at 8,000 g for 10 min to pellet particulate matter. After removal of 20 μL for protein analysis, 300 μL of supernatant for each sample was subjected to 15% (w/v) KOH hydrolysis (incubation for 1 hour at 37°C) for the measurement of total 8-isoprostanes. Samples were then neutralized by adding 600 μL of 1N HCl to each supernatant. Extraction was performed using SEP cartridge C18 columns (Thermo Fisher Scientific). Isoprostanes were eluted with methanol, dried under nitrogen, and reconstituted into 300 μL of ELISA buffer (1X). An aliquot of 50 μL of reconstituted isoprostane extract was assayed in duplicate. The result was expressed as pg/mg protein.

Malondialdehyde (MDA) levels in the placenta were measured using the thiobarbituric acid reaction (TBARS Assay, no. 10009055, Cayman Chemical Company). Briefly, 25 mg of placenta was homogenized in 250 μL of RIPA buffer containing 10 mM Tris-Cl, pH 8, 140 mM NaCl, 1% triton X-100, 0.1% sodium deoxycholate, 0.1% sodium dodecyl sulfate, 1 mM EDTA, 1 mM NaV04, 25 mM NaF, 1 mM PMSF, 1 μg/mL leupeptin, and aprotinin. Supernatants were obtained after centrifugation at 1600 x g for 10 min at 4°C. The absorbance of the supernatant fraction was read at 530 nm, and results were expressed in nmol per mg of placental tissue protein.

### 2.8. Statistical analysis

Comparisons of clinical data, placental mRNA expression, antioxidant enzymes and proteins, lipid peroxidation markers and Pearson correlations were performed using twotailed unpaired *t-*test and the association of placental histopathological features with MHO was analyzed using the Fisher’s exact test (GraphPad Prism 9.0.0, GraphPad Software, Boston, MA, USA). Differences were considered statistically significant when *p* < 0.05. Cohen’s *d* index was calculated to assess the effect size, both in the control and MHO group. Cohen’s *d* evaluates the magnitude of difference between the two groups. Values of Cohen’s *d* index below 0.19, between 0.20 and 0.49, between 0.50 and 0.79, between 0.80 and 1.29, and above 1.30 are classified as insignificant, small, average, large and very large effect, respectively [42]. For the analysis of categorical variables of placental histopathology, the effect size was evaluated using the Cramer’s V index. Values between 0.0 and 0.1, between 0.1 and 0.20, between 0.20 and 0.40, between 0.40 and 0.60, between 0.60 and 0.80, and between 0.80 and 1.0 are classified as negligeable, weak, moderate, relatively strong, strong and very strong, respectively [43].

Metabolomics data were analysed using partial least square-discriminant analysis (PLS-DA), univariate analysis (*t*-test, volcano plot), and pathway analysis in the platform MetaboAnalyst 5.0 [https://www.metaboanalyst.ca/MetaboAnalyst/upload/StatUpload-View.xhtml]. Raw data were normalized by auto scaling and then used for both univariate and multivariate analyses. The PLS-DA model was cross-validated by the evaluation of validation metrics including the predicted (Q^2^) and explained (R^2^) variances. The statistical significance of altered metabolites within experimental groups was set to false discovery rate (FDR)-corrected p-value < 0.25, raw p-value < 0.05, and fold change > 1.2 or < 0.83. The variable importance in projection (VIP) ranking analysis was then performed to investigate variable’s importance in the PLS-DA model. The VIP was used as a quantitative estimation of the discriminatory power of each individual metabolite. Metabolites with a VIP score of >1, were considered important in the PLS-DA model. Receiver operating characteristic curve (ROC) analysis was performed with SPSS 28.0 to calculate the area under the curve (AUC) to evaluate the accuracy of altered metabolites to diagnose MHO placentae. An AUC of 0.5 indicates a poor diagnostic test, while a value of 1.0 indicates an ideal test. MetaboAnalyst pathway analysis was then performed on data for all identified metabolites in tiers 1 and 2, via MetaboAnalyst 5.0, using human metabolome database (HMDB) and KEGG identifiers of metabolites, to identify the affected metabolic pathways (FDR < 0.05 and pathway impact > 0.1).

## 3. Results

### 3.1. Pregnancy and placental characteristics

Maternal and delivery characteristics of the study cohort were previously reported by Cohen et al. [16]. Mean pre-pregnancy BMI was higher in the MHO vs. control (C) group (C: 21.1 ± 2.0 kg/m^2^ vs. MHO: 42.3 ± 7.6 kg/m^2^, p < 0.0001, Cohen’s *d* = 3.8). Birth weight was significantly higher in the MHO group than the C group (C: 3170.0 ± 403.4 g vs. MHO: 3795.3 ±717.8 g, p < 0.01, Cohen’s *d* = 1.1). The characteristics of placentae of the two study groups are displayed in **Table 1**. The placenta was heavier in the MHO group (C: 592.3 ± 111.0 g vs. MHO: 762.0 ± 164.3 g, p < 0.01, Cohen’s *d* = 1.2). This was reflected in an increase in placental breadth (C: 15.1 ± 1.5 cm vs. MHO: 16.9 ± 2.0 cm), thickness (C: 1.5 ± 0.4 cm vs. MHO: 2.0 ± 0.4 cm) and surface area (C: 199.6 ± 34.9 cm^2^ vs. MHO: 243.7 ± 52.7 cm^2^) (for all measures, p < 0.05, Cohen’s *d* ≥ 1). The increase in placental length in the MHO group trended to significance (C: 16.7 ± 2.1 cm vs. MHO: 18.1 ± 2.0 cm, p = 0.068, Cohen’s *d* = 0.7). Additionally, the fetal-placental weight ratio, a measure of placental efficiency [20] [21], was not statistically different between the 2 groups (C: 5.0 ± 0.1 vs. MHO: 5.4 ± 0.2, p = 0.2821, Cohen’s *d* = 2.5). Despite the morphometric changes of the placenta, the scored placental histopathology parameters showed no significant differences between MHO and control (p > 0.999, Cramer’s < 0.3, **Table 1**).

**Table 1.** Placental morphometric and histopathological features among control (C, n=15) and metabolically healthy obesity (MHO, (n=15).

|  | C | MHO | P-value | d/V |
| --- | --- | --- | --- | --- |
| Placental weight<sup>1</sup> (g) | 592.3 ± 111.0 | 762.0 ± 164.3 | 0.0026 | 1.2 |
| Placental length (cm) | 16.7 ± 2.1 | 18.1 ± 2.0 | 0.0685 | 0.7 |
| Placental breadth (cm) | 15.1 ± 1.5 | 16.9 ± 2.0 | 0.0091 | 1.0 |
| Placental thickness (cm) | 1.5 ± 0.4 | 2.0 ± 0.4 | 0.0049 | 1.2 |
| Placental surface area (cm<sup>2</sup>) | 199.6 ± 34.9 | 243.7 ± 52.7 | 0.0116 | 1.0 |
| Distal villous hypoplasia (focal or diffuse)<sup>2</sup> | 0 (0) | 0 (0) | NS | ND |
| Syncytial knots | 0 (0) | 0 (0) | NS | ND |
| Chorangiomas | 0 (0) | 2 (13) | NS | 0.3 |
| Delayed villous maturation (focal or diffuse) | 1 (7) | 5 (33) | NS | 0.3 |
| Avascular fibrotic villi | 0 (0) | 0 (0) | NS | ND |
| Increased focal perivillous fibrin deposition | 1 (7) | 1 (7) | NS | 0.0 |
| Massive perivillous fibrin deposition | 0 (0) | 0 (0) | NS | ND |
| Maternal floor infarct pattern | 0 (0) | 0 (0) | NS | ND |
| Intervillous thrombi | 0 (0) | 0 (0) | NS | ND |
| Chronic inflammation | 0 (0) | 0 (0) | NS | ND |

<sup>1</sup>Placental morphometric data are presented as mean (±SD). BMI, body mass index. <sup>2</sup>The distributions of the different placental histopathological features are represented using frequency (n) and percentage in brackets (%). The effect size was calculated using Cohen's *d* or Cramer's *V*; NS = non-significant (*p* > 0.1); ND = not determined due to absence of observation in both groups.

### 3.2. Changes in the metabolome in MHO

With the LC–MS, 2,808 metabolite peak pairs were detected in each sample. Data within each individual placental region (centre, periphery ad middle) were separately determined and analysis conducted. There were no significant metabolite changes between MHO and control groups within these regions, allowing for pooling of each set of three placenta samples. **Figure 1** displays the pooled results of the PLS-DA analysis. A separation is observed between the two groups, with a degree of overlap between MHO and control women (**Figure 1A**). This separation is estimated by the two quality parameters, R^2^ = 0.904 for the explained variation and Q^2^ = 0.487 for the predictive capability of the model (**Figure 1B**).

**Figure 1.**
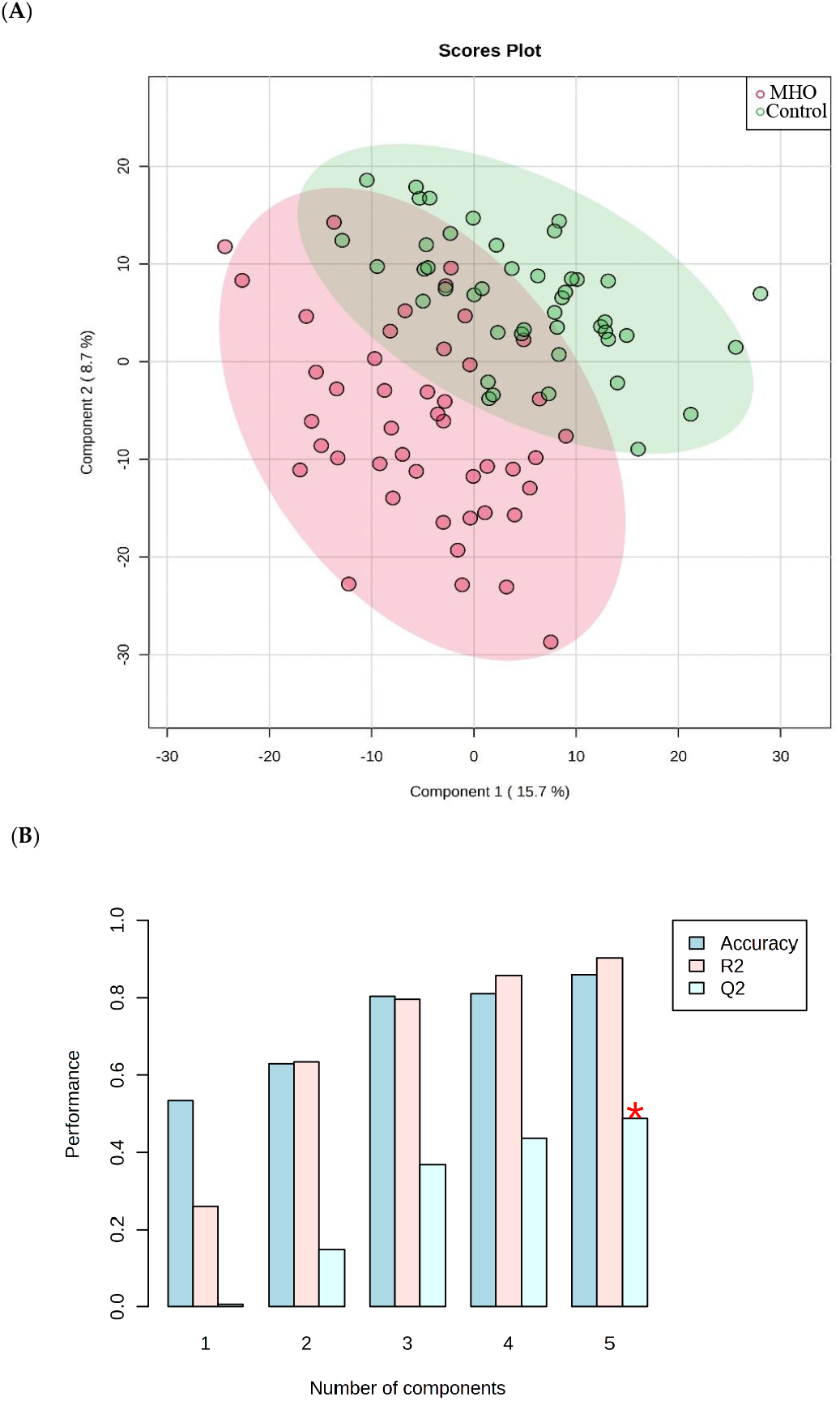
Partial least squares discriminant analysis (PLS-DA) of control and MHO groups. (**A**) The PLS-DA 2D scores showing discrimination between MHO and control groups. The red dots represent the MHO samples, and the green dots represent controls. Data from three areas of the placenta [centre, periphery (1–2 cm from the placental edge), and middle (between these two sections)] and each set of three samples per placenta was pooled. (**B**) Cross validation chart: PLS-DA classification performance showed that the best classifier was obtained using 5 components, considering the accuracy, variations (R2) and prediction of the model (Q2).

Abundance of each metabolite peak pair (intensity) was compared between MHO and control groups, using univariate analysis. Of the 2808 recorded metabolite peak pairs, 561 were successfully identified in tier 1 and tier 2 (**Supplementary Table S2**) and 37 were found to be differentially abundant between MHO and control groups (fold change > 1.2 or < 0.83, and FDR-p-value < 0.25 and raw p-value < 0.05) (**Figure 2**).

**Figure 2:**
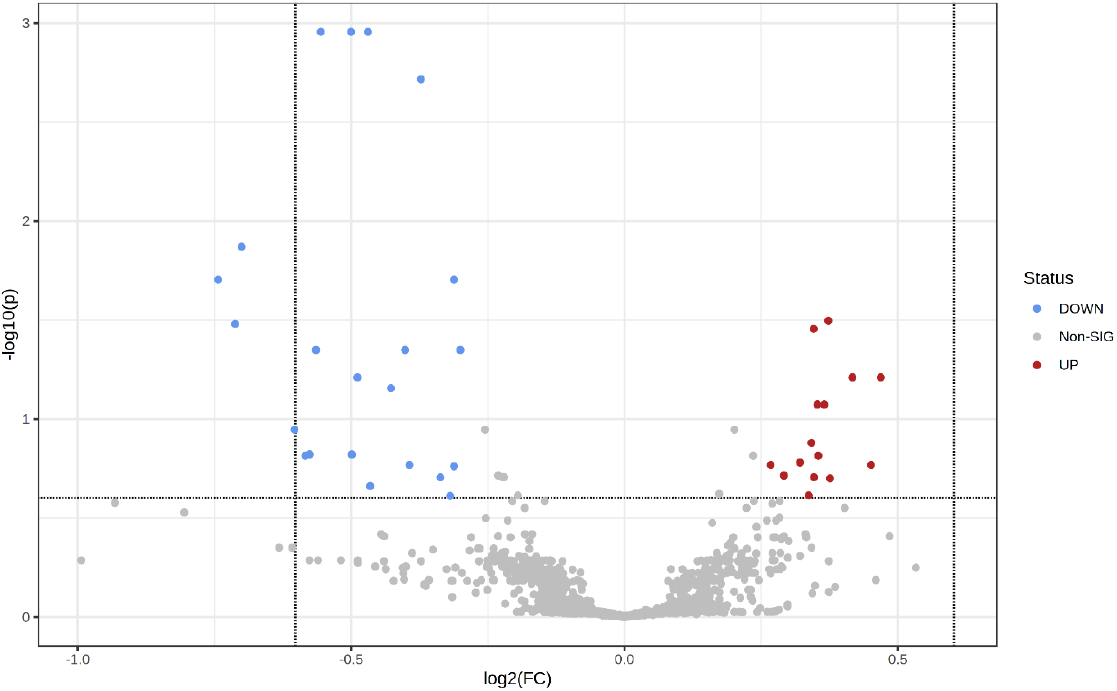
Visualization of changes in metabolite patterns by volcano plots using univariate analysis. Volcano plot shows the statistically significant altered metabolites (FDR-corrected p-value < 0.25, raw p-value < 0.05, and FC > 1.2 or < 0.83). N = 15 per each group with data from central, peripheral and middle placenta combined within each individual. The x-axis indicated log2(fold-change) vs. the MHO group, and the y-axis indicates −log10(p-value). The levels of 22 metabolites were down-regulated (blue) and 15 were up-regulated (Red) in the MHO group compared to the control group.

Among the 37 metabolite peak pairs, 22 were found with low abundance and 15 with high abundance in the MHO group. Of these differentially abundant peak pairs, 10 were successfully identified in tier 1 and tier 2 and are displayed in **Table 2**.

**Table 2.** Alterations in placental metabolites in the MHO vs. control group.

| Compound<sup>1</sup> | Fold change | p. adjusted | p. value |
| --- | --- | --- | --- |
| 3-aminoisobutanoic acid (BAIBA) | 0.6804 | 0.0011 | <0.0001 |
| 2-aminobutyric acid (2AB) | 0.7071 | 0.0011 | <0.0001 |
| Gamma-Glutamyl-Glycine (γ-Glu-Gly) | 0.8057 | 0.0198 | 0.0001 |
| Ophthalmic acid (OPT) | 0.6763 | 0.0448 | 0.0004 |
| Prolyl-Glutamine (Pro-Gln) | 0.7571 | 0.0448 | 0.0003 |
| N6-acetyl-LL-2,6-diaminoheptanedioic acid (N6-acetyl-LL-2,6-DAP) | 0.8121 | 0.0448 | 0.0003 |
| Valyl-asparagine (Val-Asn) | 0.7438 | 0.0698 | 0.0007 |
| Pyridoxal | 0.6582 | 0.1130 | 0.0015 |
| 3-hydroxykynurenamine O-sulfate |  |  |  |
| (C05636) | 0.7918 | 0.1967 | 0.0049 |
| Pyridoxal phosphate (PLP) | 0.8017 | 0.2440 | 0.0069 |

<sup>1</sup>Relative values of differentially expressed and identified metabolites in MHO placenta expressed as fold change of control placenta. Significantly different metabolites between MHO and control placenta [Fold change > 1.2 or < 0.83 and FDR-corrected p-value < 0.25 (corresponding to a raw p-value threshold of 0.049) as criteria] are presented. N = 15 per each group with data from central, peripheral and middle placenta combined within each individual placenta.

VIP scores were used to identify the metabolites contributing mostly to changes in MHO placentae (**Figure 3**). The top 50 metabolite peak pairs with the highest significance in the group discrimination (VIP > 1), identified metabolites including 3-aminoisobutanoic acid, 2-aminobutyric acid, gamma-glutamyl-glycine, ophthalmic acid, prolyl-glutamine, N6-acetyl-LL-2,6-diaminoheptanedioic acid (N6-acetyl-LL-2,6-DAP), valyl-asparagine, pyridoxal, 3-hydroxykynurenamine O-sulfate, and pyridoxal phosphate that were downregulated in MHO. These had VIP scores > 2 and were considered relevant metabolites.

**Figure 3:**
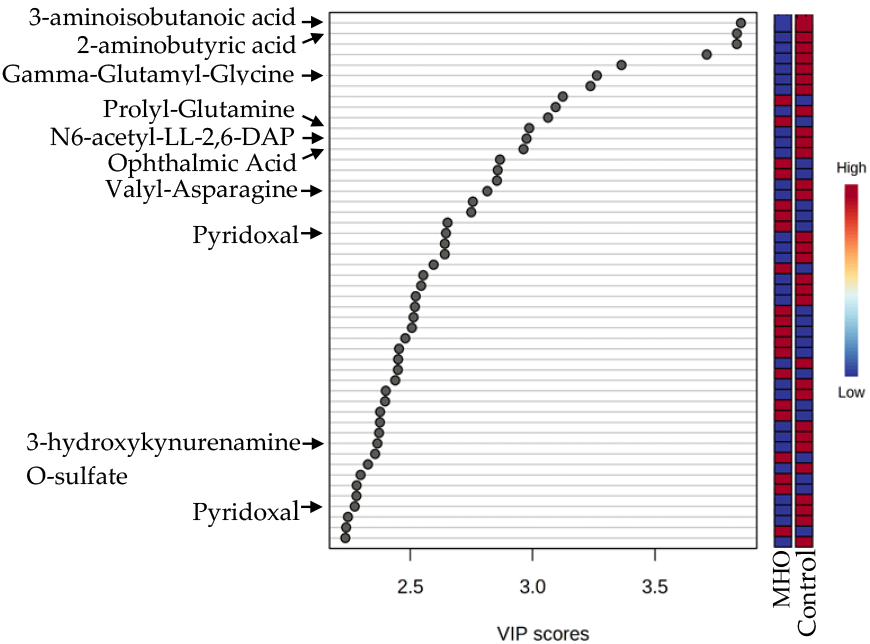
Variable importance in projection (VIP) scores of the PLS-DA analysis. VIP scores indicate the top 50 metabolites contributing to the separation of metabolic profiles in control vs. MHO groups. The relative abundance of metabolites is indicated by a colored scale from blue to red representing the low and high, respectively. Identified metabolites with a high confidence are displayed in the left of the chart.

ROC curve analysis revealed the diagnostic accuracy of these metabolites in discriminating between the control and MHO group. All identified metabolites, with the exception of 3-hydroxykynurenamine and pyridoxal phosphate, individually showed a high diagnostic accuracy (AUC ≥ 0.7, p < 0.01) (**Figure 4A**). The diagnostic accuracy of individual top-ranked metabolites based on VIP scores including 3-aminoisobutanoic acid, 2-aminobutyric acid, gamma-glutamyl-glycine and N6-acetyl-LL-2,6-diaminoheptanedioic was lower than the curve obtained from their combination (AUC = 0.831, p < 0.0001) (**Figure 4B**). Similar observations were made for metabolites related to oxidative stress, including 2-aminobutyric acid, ophthalmic acid, pyridoxal and pyridoxal phosphate (AUC = 0.788, p < 0.0001) (**Figure 4B**).

**Figure 4:**
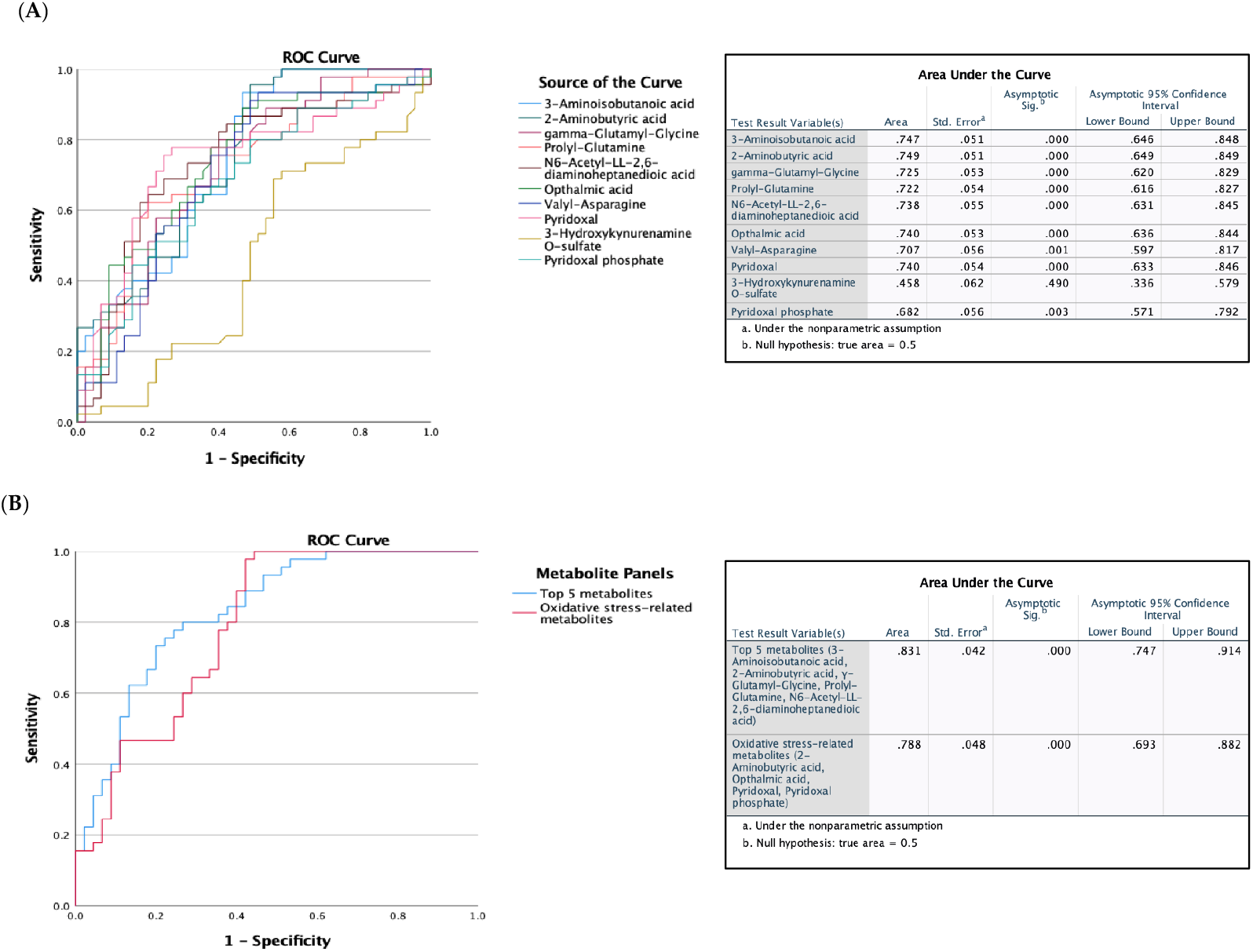
Receiver operating characteristic (ROC) curves constructed with the logistic regression model and area under the curve (AUC) tables for discrimination between control and MHO placentae. (**A**) ROC curves and AUC table showing the accuracy of individual metabolites to diagnose MHO placentae. (**B**) ROC curves and AUC table showing the accuracy of combined metabolites to diagnose MHO placentae.

A graphical summary of the identified pathways and their relative impact is shown in **Figure 5**. Overall, the perturbed pathways closely related to amino acid and vitamin B6 metabolism, and included cysteine and methionine metabolism, alanine, aspartate and glutamate metabolism, phenylalanine, tyrosine, and tryptophan biosynthesis, phenylalanine metabolism, glyoxylate and dicarboxylate metabolism, lysine degradation, amino-acyl-tRNA biosynthesis, and arginine biosynthesis (FDR < 0.05; impact values > 0.1).

**Figure 5:**
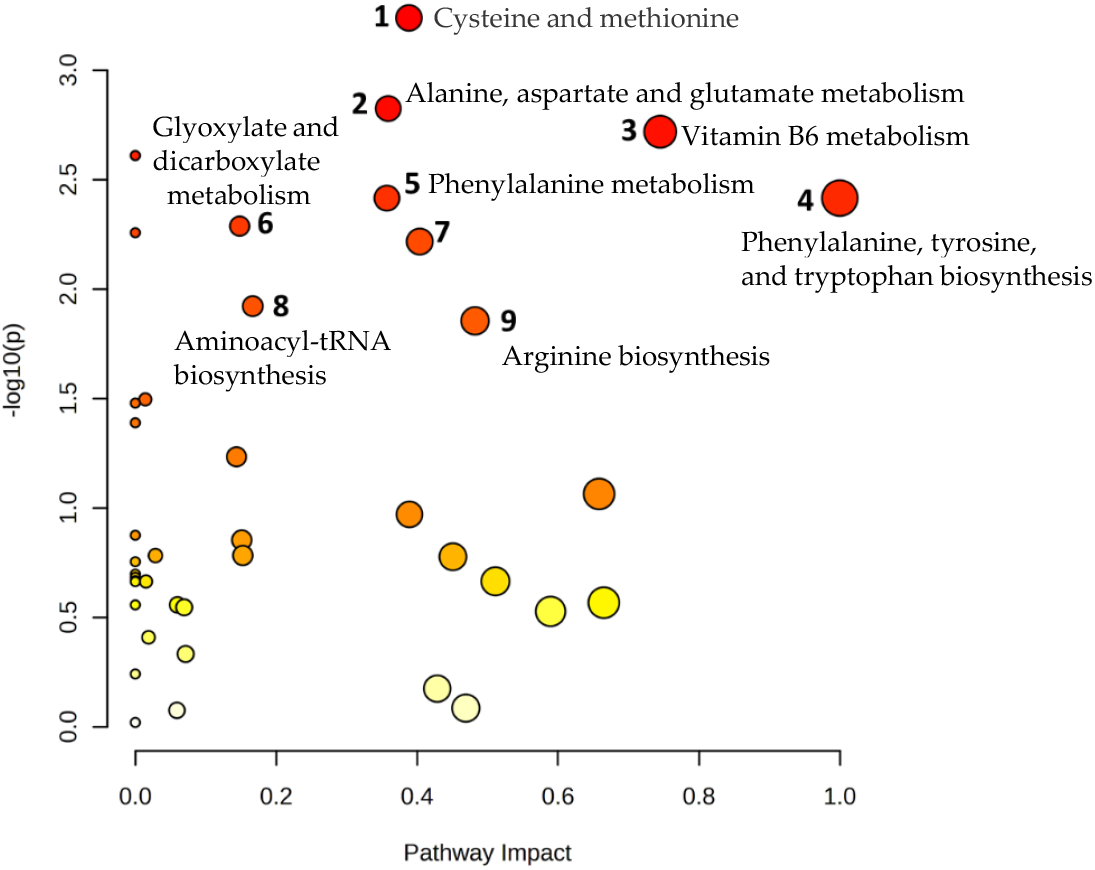
Pathway analysis by MetaboAnalyst. X-axis shows pathway impact values from pathway topology analysis; y-axis displays matched pathways from pathway enrichment analysis arranged by −log(p-value). Red indicates the most significant effects according to p value and the node size is determined by pathway impact value. The 9 altered pathways (FDR < 0.05 and pathway impact > 0.1) are displayed.

Significant downregulation of the glutathione analog ophthalmic acid and its precursor 2-aminobutyric acid were captured in the top altered metabolic pathway, the cysteine and methionine metabolism pathway (**Figure 6**).

**Figure 6:**
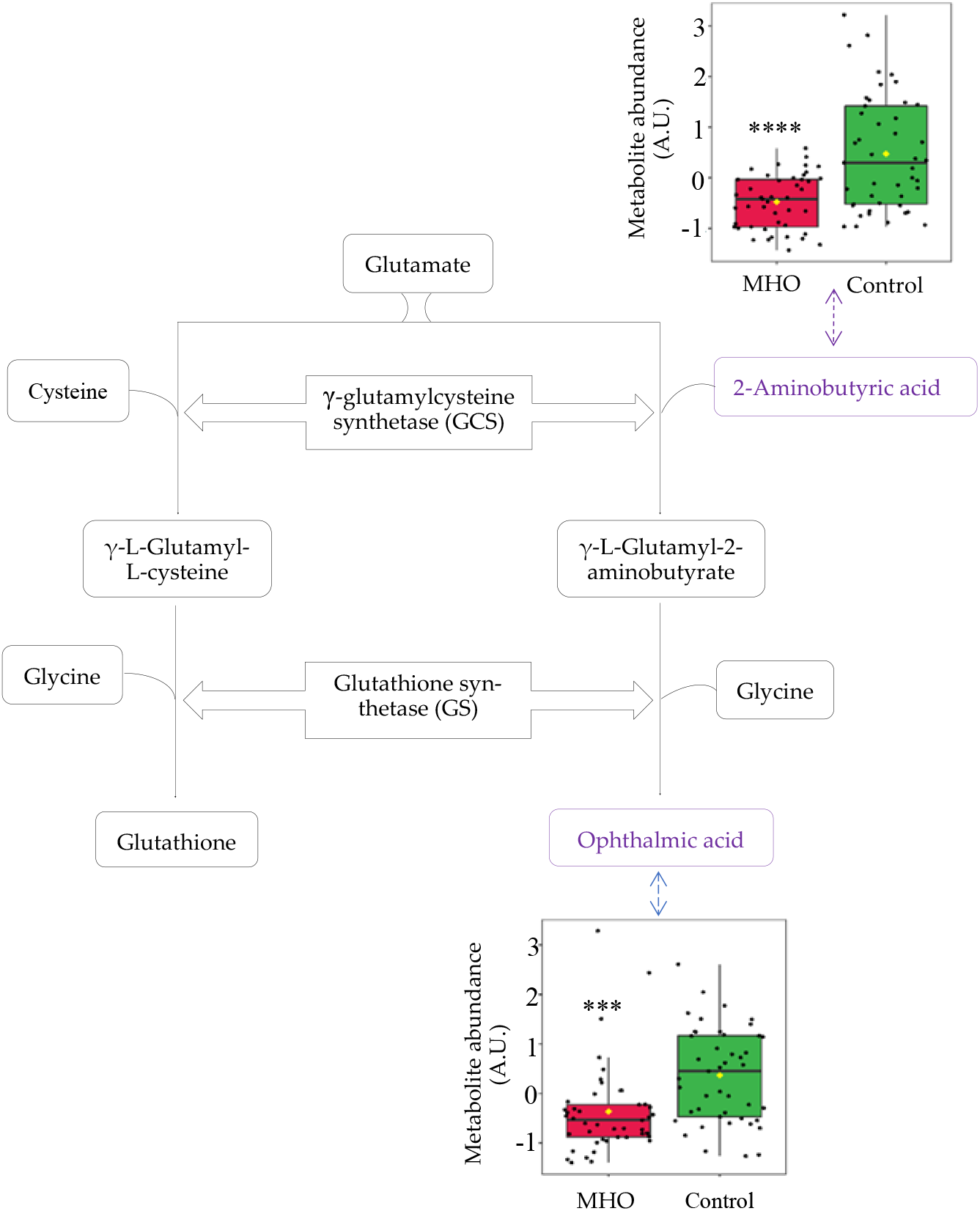
Observed metabolite changes mapped onto a maintained glutathione and reduced ophthalmic acid biosynthesis in MHO placentae. Box plots of 2-aminobutyric acid and ophthalmic acid abundance (metabolite peak pair intensity) in MHO (red boxes) and control (green boxes) groups. **** and *** Significantly different between MHO and control placentas, p-value < 0.0001 and 0.001, respectively. N = 15 per group.

Significant decrease in pyridoxal and pyridoxal phosphate, known to be involved in defense against cellular oxidative stress, were highly responsible for vitamin B6 metabolism pathway alteration in the MHO group (**Figure 7**).

**Figure 7:**
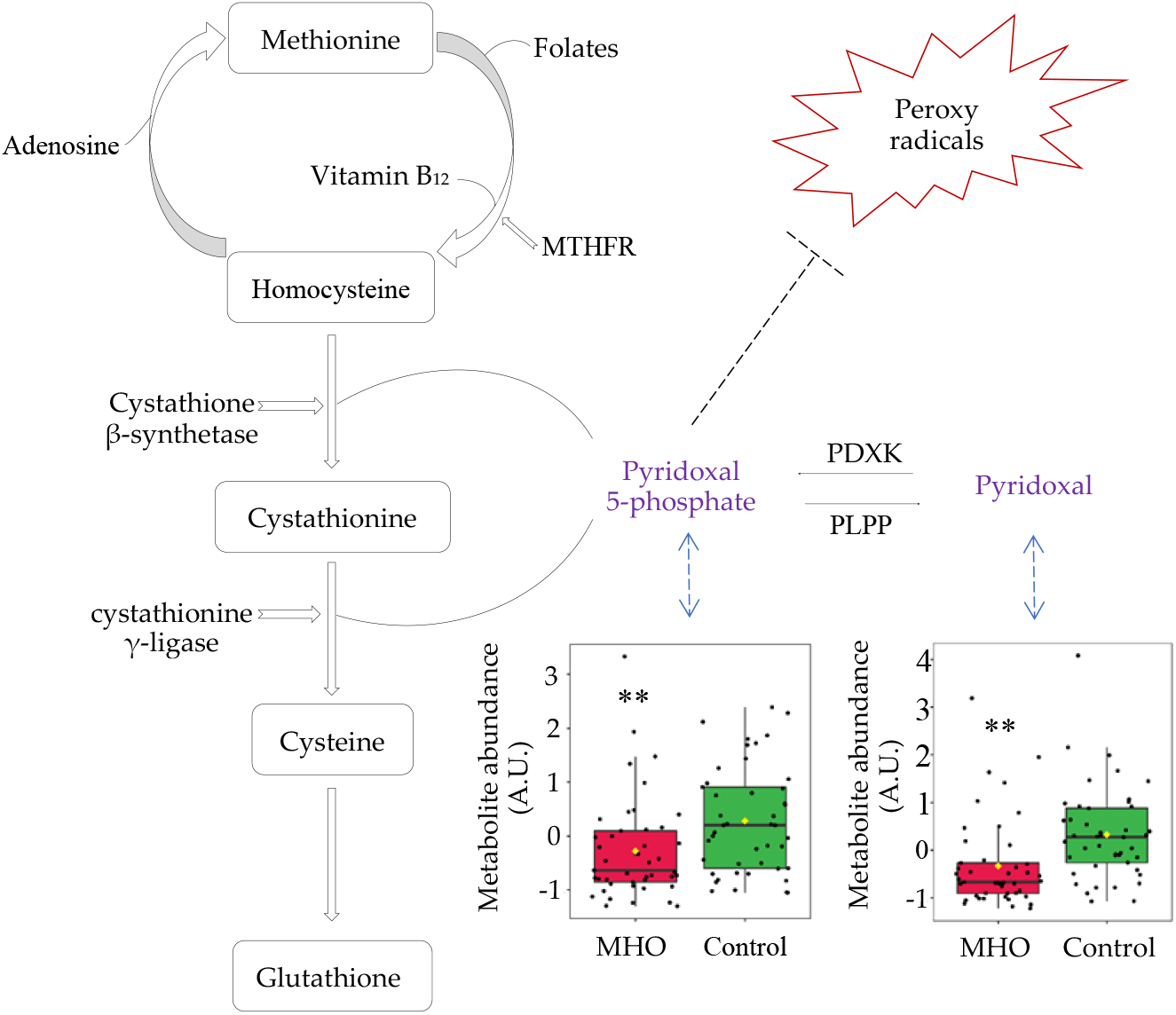
Metabolites changes revealed decrease in vitamin B6 forms in MHO placentae. Pyridoxal phosphate (pyridoxal 5’-phosphate) is the active form of vitamin B6 (pyridoxine or pyridoxal) and acts as a coenzyme in the transulfuration and glutathione biosynthesis. Pyridoxal phosphate may also directly react with the peroxy radicals and thereby scavenge radicals. Box plots of pyridoxal and pyridoxal phosphate abundance (metabolite peak pair intensity) in MHO (red boxes) and control (green boxes) groups are displayed. ** Significantly different between MHO and control placentae, *p* < 0.01. N = 15 per group. MTHFR = methionine synthase reductase, PDXK = pyridoxal kinase, PLPP = pyridoxal 5-phosphate phosphatase.

### 3.3. Relationship between metabolite changes and placental measurements

To understand the relation between metabolite changes and placental characteristics, linear correlations were performed with metabolite abundance in the 2 study groups and placental weight, thickness, breadth and surface area (**Table 3**). Reduced abundance of 3-aminoisobutanoic acid and 2-aminobutyric acid were correlated with increased placental weight, thickness and breadth (p < 0.01). In addition, reduced abundance of gamma-glutamyl-glycine and ophthalmic acid were only correlated with increased placental thickness and breadth (p < 0.05). The low abundance of N6-acetyl-LL-2,6-DAP was however correlated with increased placental weight and thickness. Further, abundance of valyl-asparagine and pyridoxal decreased while placental weight and breadth increased (p ≤ 0.05). Finally, placental thickness and weight increased while abundance of prolyl-glutamine and pyridoxal phosphate decreased, respectively (p < 0.05).

**Table 3:** Pearson correlation (r) and *p* values of linear regression for each metabolite with placental weight, thickness, breadth and surface area.

| Metabolite | Placental weight |  | Placental thickness |  | Placental breadth |  | Placental surface area |  |
| --- | --- | --- | --- | --- | --- | --- | --- | --- |
| | $r$ | $p$ | $r$ | $p$ | $r$ | $p$ | | |
| BAIBA | -0.2961 | 0.0046* | -0.5406 | <0.0001* | -0.2792 | 0.0077* | 0.1339 | 0.2083 |
| 2-aminobutyric acid | -0.2927 | 0.0051* | -0.5379 | <0.0010* | -0.2754 | 0.0086* | -0.1359 | 0.2014 |
| γ-Glu-Gly | -0.0939 | 0.3787 | -0.3125 | 0.0075* | -0.2448 | 0.0200* | -0.1715 | 0.1060 |
| Ophthalmic acid | -0.1531 | 0.1497 | -0.2563 | 0.0298* | -0.2283 | 0.0304* | -0.1709 | 0.1073 |
| Prolyl-glutamine | -0.1345 | 0.2061 | -0.2930 | 0.0125* | -0.1086 | 0.2061 | -0.0696 | 0.5144 |
| N6-acetyl-LL-2,6-DAP | -0.3018 | 0.0038* | -0.4134 | 0.0003* | -0.1559 | 0.1423 | -0.0959 | 0.3688 |
| Valyl-asparagine | -0.2324 | 0.0540* | -0.3960 | 0.1569 | -0.1282 | 0.0164* | -0.1098 | 0.3030 |
| Pyridoxal | -0.3432 | 0.0009* | -0.2125 | 0.0731 | -0.2640 | 0.0119* | -0.1826 | 0.0849 |
| C05636 | -0.0625 | 0.5583 | -0.0768 | 0.5211 | -0.0168 | 0.8752 | -0.0701 | 0.5115 |
| Pyridoxal phosphate | -0.2298 | 0.0293* | -0.2265 | 0.5580 | -0.0720 | 0.4998 | 0.0406 | 0.7041 |

\*Statistical significance ( $p < 0.05$ ); $N = 15$ per each group (MHO vs. control groups). BAIBA = 3-aminoisobutanoic acid, $\gamma$ -Glu-Gly = gamma-glutamyl-glycine, N6-acetyl-LL-2,6-DAP = N6-acetyl-LL-2,6-diaminohexanedioic acid and C05636 = 3-hydroxykynurenamine O-sulfate.

### 3.4. Placental antioxidant defense and proinflammatory markers

*Superoxide dismutase 1* (*SOD1*), *catalase* (*CAT*), *glutathione peroxidase* (*GPX*), *Glutathione synthetase* (GSS) and *glutamate-cysteine ligase modifier subunit* (*GCLM*) mRNA expression were not affected in MHO versus control groups (p > 0.10, Cohen’s *d* **≤** 0.3, **Figure 8A**). Conversely, a moderate decreased expression of *SOD 2* in the MHO group (−21%, p = 0.0594, Cohen’s *d =* 0.7) was observed. The mRNA expression of proinflammatory markers including *tumor necrosis factor α* (*TNFα*), *interleukin 6, 10* (*IL6, IL10*), *monocyte chemoattractant protein 1* (*MCP1*) and *toll-like receptor 3* (*TLR3*) was not affected in the MHO group (p > 0.10, Cohen’s *d* **<** 0.3, **Figure 8A**). While SOD activity was not significantly impacted (p = 0.10, Cohen’s *d* = 0.6), CAT activity was lower in the MHO group (−16%, p < 0.05, Cohen’s *d* **=** 0.7) (**Figures 8B, C**). GSH and GSSG concentrations and ratio of GSH:GSSG were not different between MHO and control groups (p > 0.7, Cohen’s *d* **<** 0.2; **Figures 8 D, E** and **F**).

**Figure 8:**
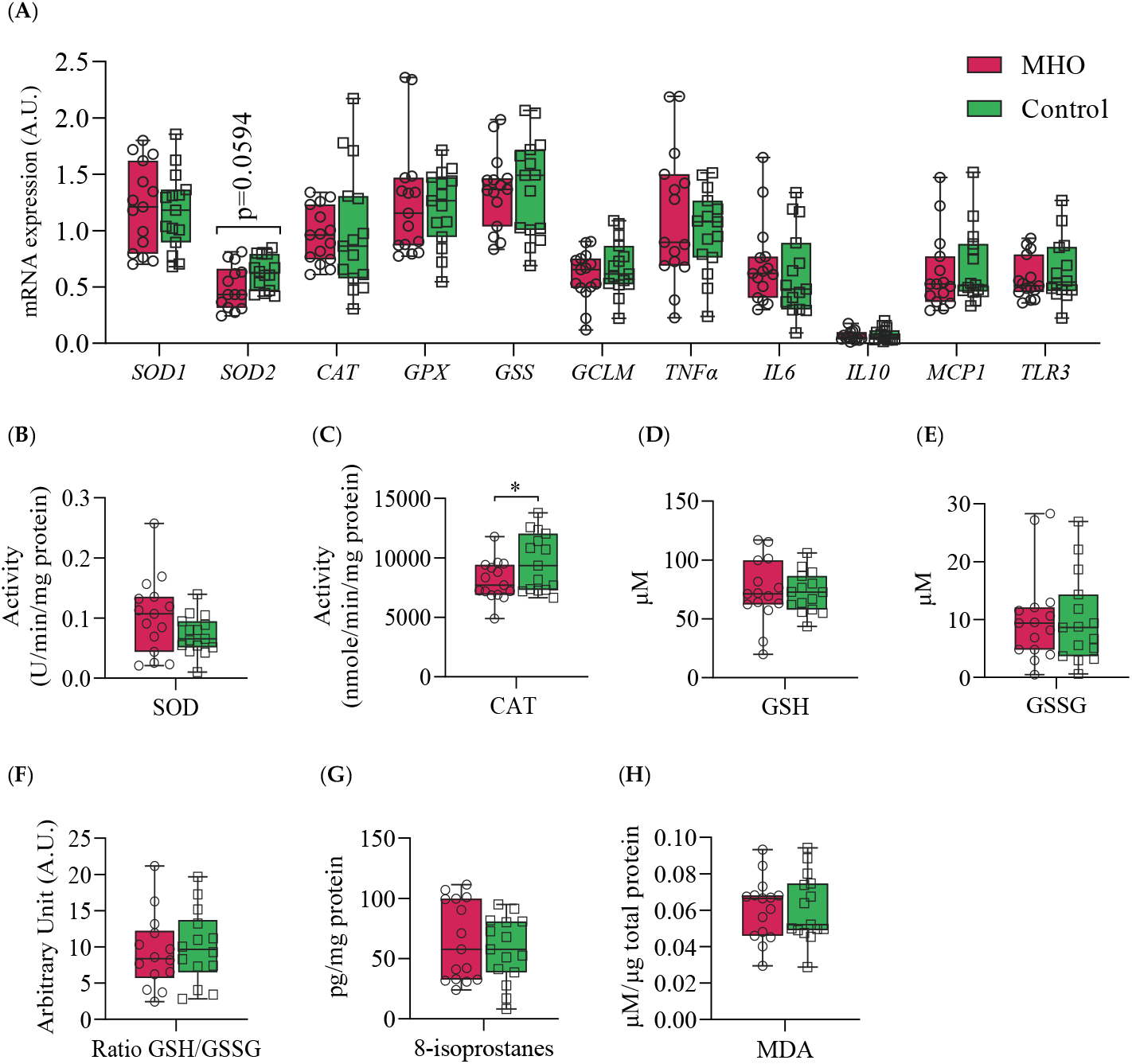
Antioxidant and proinflammatory genes, antioxidant enzymes and proteins, and lipid peroxidation markers. Panel (**A)** shows mRNA expression of *superoxide dismutase 1 & 2* (*SOD1 & 2*), *catalase* (*CAT*), *glutathione* (*GPX*), *glutathione synthetase* (*GSS*) and *glutamate-cysteine ligase modifier subunit* (*GCLM*) and inflammatory markers *tumor necrosis factor α* (*TNFα*), *interleukin 6* (*IL6*), *interleukin 10* (*IL10*), *monocyte chemoattractant protein 1* (*MCP1*) and *Toll-like receptor 3* (*TLR3*). Panels (**B)** and (**C)** indicate placental SOD and CAT activities, respectively. Panels (**D, E** and **F)** show reduced glutathione (GSH), oxidized GSH (GSSG), and GSH/GSSG ratio, respectively. Panels (**G)** and (**H)** display placental total 8-isoprostanes and Malondialdehyde (MDA) levels, respectively. Data are box plots with min to max of 14-15 individuals per group. *, indicates *p* < 0.05 when comparing the MHO *vs*. control group.

### 3.5. Lipid peroxidation markers

The two universal parameters related to the development of oxidative stress, 8-isoprostanes and malondialdehyde (MDA) levels, hallmark of arachidonic acid and lipid peroxidation respectively, showed no significant changes in MHO versus control groups (p > 0.5, Cohen’s *d* **<** 0.3; **Figures 8 G, H**).

## 4. Discussion

The current study’s primary objective was to determine whether MHO pregnancies—lacking cardiometabolic comorbidities—exhibit distinct placental morphometry and metabolomic profiles compared to normal-weight pregnancies. The findings indicated that uncomplicated pregnancies with maternal obesity classified as MHO pregnancies were associated with large, thick, and heavy placentae. Additionally, although there were no significant changes in placental histopathology, multiple metabolites were altered, and metabolic pathways were impacted. The metabolite 3-aminoisobutanoic acid was the top-ranked metabolite in differentiating MHO and control placentae, and the cysteine and methionine and vitamin B6 metabolism pathways were among the most distinct signatures for MHO placentae. The biosynthesis of 2-aminobutyric acid, which serves as a marker of glutathione depletion [44], as well as the antioxidant forms of vitamin B6, pyridoxal and pyridoxal phosphate, were reduced in MHO placentae [45,46]. This occurred alongside a decline in the activity of the antioxidant enzyme catalase; however, superoxide dismutase activity, glutathione levels, and lipid peroxidation markers remained consistent between control and MHO pregnancies. Despite the unchanged levels of superoxide dismutase, glutathione, and lipid peroxidation markers, this selective decrease in catalase activity suggests a nuanced shift in the antioxidant system that may be specific to MHO. These observations indicate that, alongside alterations in placental morphometric characteristics, there exists a specific metabolomic signature showing a partial decline in the antioxidant response, without evidence of oxidative or tissue damage in the MHO placenta.

The placenta plays a crucial role in intrauterine programming, and its phenotype has been shown to serve as a more effective indicator of disease risk later in life than other identifiers of exposure to suboptimal conditions during intrauterine development [47]. Recent findings have revealed several links between placental morphometric features, such as weight, length, width, and thickness, with diseases early and later in life [48,49]. These gross morphometric characteristics are considered biologically linked to the functional capacity of the placenta, or they may impose mechanical or other constraints on the developing fetus [48]. Moreover, changes in placental morphometric characteristics are commonly associated with maternal obesity [27,50]. However, the association between placental morphometric characteristics and maternal MHO during pregnancy is limited [27], even though women classified as having MHO constitute the majority of obese pregnancies in reported studies [13]. Although maternal obesity has been linked to placental overgrowth, relatively few studies focus on the subset of women with MHO. The finding that MHO pregnancies show a significant increase in placental thickness—without the metabolic disturbances common to MUO—underscores the importance of distinguishing these subtypes. In the current study, a significant increase in placental thickness was observed in MHO pregnancies, consistent with previous findings in uncomplicated pregnancies as reported by Bianchi et al. [27]. Our findings, however, are in contrast with this study [27], in which unchanged placental weight, diameter and area were reported in pregnancies with obesity and without GDM. However, the study by Bianchi et al. reported significant lower gestational weight gain in pregnancies with obesity, which was not observed in our cohort [16]. The findings of the present study emphasize specific placental morphometric changes, suggesting that these features may be unique to obesity and not necessarily associated with conditions like GDM, thereby underscoring the importance of accounting for BMI in studies that assess pregnancy complications where obesity is a significant risk factor.

In line with an increased placental breadth and surface area, the MHO pregnancies also displayed increased birth weights. Placental breadth is associated with neonatal body size and the size of the breadth is an indicator of the transfer of nutrients from mother to fetus; the larger the breadth the greater the transfer, although underlying processes are unknown [23]. Additionally, studies have shown that placental surface area is proportional to the number of uterine spiral arteries available for maternal blood supply, and thickness reflects the level of villous ramification and is strongly associated with fetal growth parameters [25,26]. Placental ratios including birth weight/placental weight and birth weight/placental volume have been suggested to predict pregnancy complications such as GDM and preeclampsia before delivery [22]. It is interesting to note that birth weight/placental ratio was unchanged in the MHO pregnancies, as previously reported in obese pregnancy without GDM [27]. This difference possibly underscores the uncomplicated nature of the studied pregnancies, unlike obese + GDM pregnancies in which the fetal/placental ratio is reduced [27]. The observed data, combined with placental morphometric changes, suggest that there is a compensatory and/or enhanced placental growth [51], to accommodate increased metabolic demands arising from accelerated fetal growth in these pregnancies. Such adaptations in placental growth could serve as an early compensatory mechanism, potentially influencing fetal nutrient transfer and posing long-term metabolic or cardiovascular consequences for the offspring.

Whereas placental morphometric readouts afford us one measure of placental growth as a measure of the in utero environment, placental metabolomics provides a comprehensive understanding of placental function and subsequent metabolism. In the current study, a VIP plot of metabolomics analysis highlighted that 3-aminoisobutanoic acid is the most crucial metabolite that is different between MHO and control placentae. This metabolite, also known as 3-aminoisobutyrate, is part of the beta-amino acids and derivatives class of organic compounds and is produced from ureidoisobutyric acid. It appears to be involved in stimulating free fatty acid oxidation and glucose uptake in metabolically active tissues such as adipose tissue, liver, and skeletal muscle [52]. Previously reported low levels of circulating 3-aminoisobutanoic acid found in obesity [53] align with the findings of the current study, which showed that 3-aminoisobutanoic acid was reduced in MHO placentae and inversely correlated with placental weight, thickness, and breadth. These data imply that 3-aminoisobutanoic acid could be a potential metabolic biomarker for enhanced placental growth in MHO pregnancies, and its depletion may contribute to accelerated fetal growth, impacting the lifelong health of the offspring in MHO pregnancies. As highlighted, 3-aminoisobutanoic acid surfaced as a leading candidate biomarker for MHO-related placental adaptations. Its inverse correlation with placental dimensions emphasises its potential utility in predicting or diagnosing MHO at earlier gestational stages.

In the current study, reductions in ophthalmic acid and 2-aminobutyric acid, metabolites in the cysteine and methionine metabolism pathway, were among the most distinct signatures for MHO pregnancy. Additionally, ophthalmic acid and 2-aminobutyric acid were inversely correlated with placental weight, thickness and breadth. Ophthalmic acid, also known as ophthalmate, is synthesised *in vivo* from 2-aminobutyric acid through the same enzymatic machinery as glutathione, a key antioxidant tripeptide and one of the most abundant intracellular antioxidants [54]. Interestingly, ophthalmic acid lacks a reducing cysteine moiety, and therefore, it is far more stable than glutathione [55,56]. Taken together, the altered cysteine and methionine metabolism pathway and the lack of changes in both reduced and oxidised glutathione indicate a preserved glutathione response in MHO placentae. This is in accordance with previous reports of a placental adaptation of the affected antioxidant response towards a nitric oxide-induced alternative pathway and highlights changes in the reactive oxygen species/reactive nitrogen balance in order to reduce oxidative damage and preserve placental function in MHO pregnancy [15].

Another antioxidant system that is impacted in MHO placentae is the vitamin B6 pathway, which is a central pathway required for many processes ranging from amino acid biosynthesis and catabolism, fatty acid biosynthesis and breakdown, antioxidant, as well as the biosynthesis of neurotransmitters [45,46,57]. The metabolites pyridoxal and pyridoxal phosphate (PLP) are among the major forms of vitamin B6 [58], and within cells, PLP is the active B6 vitamin [59]. In the present study, significant disturbances in the vitamin B6 metabolism pathway, including reduced pyridoxal and PLP, were observed in MHO placentae. Bjørke-Monsen et al. have demonstrated that pre-pregnancy BMI is inversely related to PLP [60], and postulated that lower pyridoxal and PLP levels may contribute to adverse pregnancy outcomes associated with maternal obesity, as an optimal micronutrient status is vital for normal fetal development[60]. In this sense, our data show that pyridoxal and PLP were both inversely related to placental weight and breath in MHO placentae.

Vitamin B6 also plays a significant antioxidant role in preventing the decrease of the antioxidant enzyme catalase in peripheral tissues *in vivo and in vitro*, through scavenging superoxide radicals [45,46]. Interestingly, parallel to lower levels of B6 isoforms, a decrease in the activity of the antioxidant enzyme catalase in MHO placentae is observed. While these indicators suggest a potential early failure in the placental antioxidant defense system, protective mechanisms seem to be still active as there were no changes in malondialdehyde (MDA) and 8-isoprostane, oxidative stress markers formed by the peroxidation of lipid and arachidonic acid, respectively [61,62]. These data, combined with the alterations in glutathione analogs (ophthalmic acid and 2-aminobutyric acid) and the absence of changes in superoxide dismutase activity, indicate a partial decline in the antioxidant response without oxidative damage in MHO placentae. This is in agreement with reports of decreased activity of both catalase and superoxide dismutase antioxidant enzymes and no significant change in the levels of MDA in full-term placentae from MHO pregnancies [15].

Inflammation is known to be associated with placental dysfunction and pregnancy complications [63]. Elevated levels of inflammatory markers, such as interleukin 6 (IL-6), IL-8, IL-1β, and monocyte chemotactic protein-1 (MCP-1), are observed in both maternal plasma and the placenta in cases of maternal obesity [64]. This cytokine profile is driven by several inflammatory pathways including activation of Toll-like receptor 4 (TLR4) and the promotion of nuclear factor kappa light chain enhancer of activated B cells (NF-κB), leading to increased generation of reactive oxygen species and the secretion of proinflammatory cytokines such as IL-6, IL-8, MCP-1, tumor necrosis factor α (TNF-α), and IL-1β [64]. In the current study, no significant changes in mRNA expression of proinflammatory markers *TNFα, IL6, IL10, MCP1* and *TLR3* were observed in MHO placentae. In contrast to MUO pregnancies, where inflammation is typically heightened [64], the absence of significant changes in these proinflammatory transcripts in MHO further suggests that metabolic health status—rather than BMI alone—may shape the placental inflammatory milieu.

The evaluation of Receiver Operating Characteristic (ROC) curves is a standard method for comparing various predictors across a range of values. Beyond visual estimation, the calculation of the area under the curve and the application of test algorithms aid in identifying the strongest predictor [65]. Using the current dataset, 3-aminoisobutanoic acid, 2-aminobutyric acid, Gamma-Glutamyl-Glycine, Prolyl-Glutamine, N6-acetyl-LL-2,6-DAP, Ophthalmic Acid, Valyl-Asparagine, and Pyridoxal metabolites were effective in identifying MHO placentae with AUC values greater than or equal to 0.7. Notably, O-sulfate 3-aminoisobutanoic acid, 2-aminobutyric acid, prolyl-glutamine, and N6-acetyl-LL-2,6-diaminoheptanedioic or N6-acetyl-LL-2,6-DAP acid, along with Valyl-asparagine, when combined, formed an outstanding group marker for MHO placentae with AUC values of 0.8. Future development of predictive assays that incorporate these metabolites— either individually or in combination—could facilitate early identification of MHO pregnancies, thereby enabling more tailored clinical management approaches.

The selection of study participants with no clinical diagnosis of pre-existing diseases or obesity-related pregnancy complications, all of whom underwent caesarean deliveries, ensures that the cohort represents pregnancies without cardiometabolic comorbidities. Moreover, standardising the mode of delivery was intended to avoid labor-induced metabolic fluctuations and provide a clearer assessment of placental metabolites. However, it is important to note certain limitations. Although including only caesarean deliveries controlled for labor-induced metabolic and inflammatory changes, future research should include placentae from both vaginal and elective caesarean section births. Maternal blood samples obtained both at term and postpartum would be helpful in assessing whether these findings hold in broader obstetric contexts. Furthermore, while sample group stratification based on sex is critically important to report, due to the relatively small sample size of the MHO and control groups, fetal sex-based analyses was not able to be performed. In addition, in this preliminary study, the impact of maternal MUO has not been evaluated. The spectrum of obesity is generally considered a continuum with MHO eventually leading to MUO, and our MHO pregnancies may have been subclinical in their progress to MUO. Additionally, incorporating the analysis of supplementary matrices like plasma samples will be crucial to further enhance the clinical applicability of these findings. Finally, integrating the assessment of plasma triglycerides and whole-body insulin sensitivity into the inclusion criteria for MHO [66] will strengthen our classification of MHO pregnancies. Nonetheless, this study identifies morphometric and metabolic differences between MHO and normal weight placentae, providing valuable preliminary data to inform the development of new hypotheses.

## 5. Conclusions

Building on the theme of altered placental morphology and metabolomic profiles in uncomplicated metabolically healthy obese (MHO) pregnancies, this study highlights how metabolomic profiling can aid in identifying pregnancy-related biomarkers, assessing placental and fetal well-being, predicting outcomes, and promoting personalised prenatal care [67]. Changes in several biochemical pathways, including cysteine and methionine and vitamin B6 metabolism, were among the most distinct signatures for MHO pregnancy. These metabolic changes underscore the practical value of assessing maternal metabolic health and their potential influence on the lifelong health of offspring. Nevertheless, further validation through larger cohort studies and broader biomarker sampling is warranted. Moreover, the reported placental morphometric and metabolomic changes collectively lay the groundwork for future research on screening and managing metabolically healthy obese (MHO) pregnancies, alongside maternal and offspring health outcomes. These findings emphasize the importance of controlling for the effects of obesity itself when studying metabolic markers in obesity-related pregnancy complications. Focusing on health markers, rather than relying solely on weight or BMI, is essential to fully capture the nuanced impact of maternal obesity on placental function and fetal development. A clear distinction between MHO and MUO, as well as metabolically healthy and unhealthy states in non-obese populations, remains vital for advancing both research and clinical management strategies.

## Supplementary Materials

The following supporting information can be downloaded at: www.mdpi.com/xxx/s1, Table S1: Primer sequences used to measure mRNA expressions by RTqPCR; Table S2: List of peak pairs detected from CIL LC-MS measurement of the samples (experimental data with imputed value).

## Author Contributions

Conceptualization, O.S., G.E., B.de V., and T.R.H.R.; methodology, O.S., A.R., S.Z., X.W., and D.G; software, O.S., A.R., S.Z., and X.W.; validation, O.S., A.R., S.Z., X.W., and D.G; formal analysis, O.S., A.R., S.Z., X.W., and D.G; investigation, O.S., A.R., S.Z., X.W., D.G., G.E., L.L., B.de V., and T.R.H.R.; resources, G.E.,., B.de V., L.L., and T.R.H.R.; data curation, O.S., A.R., S.Z., and X.W.; writing—original draft preparation, O.S.; writing—review and editing, O.S., G.E., B.de V., and T.R.H.R.; visualization, O.S. and A.R; supervision, B.de V., and T.R.H.R; project administration, B.de V.; funding acquisition, G.E., and B.de V. All authors have read and agreed to the published version of the manuscript.

## Institutional Review Board Statement

The study was conducted in accordance with the Declaration of Helsinki and approved by the Western University’s Human Subjects Research Ethics Board (REB# 106663).

## Informed Consent Statement

Informed consent was obtained from all subjects involved in the study.

## Disclaimer/Publisher’s Note

The statements, opinions and data contained in all publications are solely those of the individual author(s) and contributor(s) and not of MDPI and/or the editor(s). MDPI and/or the editor(s) disclaim responsibility for any injury to people or property resulting from any ideas, methods, instructions or products referred to in the content.

