## Supplementary Table S1 for "Altered placental morphology and metabolomic profile in association with uncomplicated metabolically healthy obese pregnancy"

**Table S1:** Primer sequences used to measure mRNA expressions by RTq‐PCR.

| **Gene** | **Accession**  **number** | **Annealing**  **temperature** | **Forward/reverse primer** |
| --- | --- | --- | --- |
| *SOD1* | NM­ˍ000454.5 | 59 ºC | Forward: 5'-GGGGAAGCATTAAAGGACTG-3'  Reverse: 5'-CCACCGTGTTTTCTGGATAG-3' |
| *SOD2* | NMˍ001322820.2 | 62 ºC | Forward: 5'- GCGTTTACTCTTAGCAGAAGCTC-3'  Reverse: 5'- GGTGACGTTCAGGTTGTTCA-3' |
| *CAT* | NMˍ001752.4 | 64 ºC | Forward: 5'-GCCACAGGAAAGTACCCCTC-3'  Reverse: 5'- CGGTGAGTGTCAGGATAGGC-3' |
| *GPX* | NMˍ001329503.2 | 64 ºC | Forward: 5'-ACACCCAGATGAACGAGCTG-3'  Reverse: 5'-CAAACTGGTTGCACGGGAAG-3' |
| *GSS* | NMˍ000178.4 | 60 ºC | Forward: 5'-GGGGTATCCTCCTAAAGACC-3'  Reverse: 5'-CACTCAGTCCTATCCCAAGT-3' |
| *GCLM* | XMˍ047418031.1 | 60 ºC | Forward: 5'-GAAGAAGTGCCCGTCCA-3'  Reverse: 5'-GGTGAAGCAATGATCACAGA-3' |
| *TNFα* | NMˍ000594.4 | 60 ºC | Forward: 5'-AAACGGAGCTGAACAATAGG-3'  Reverse: 5'-ATTACAGACACAACTCCCCT-3' |
| *IL6* | NMˍ000600.5 | 60 ºC | Forward: 5'-CTTCGGTCCAGTTGCCTT-3'  Reverse: 5'-CCATCTTTGGAAGGTTCAGG-3' |
| *IL10* | NMˍ000572.3 | 62 ºC | Forward: 5'- TCTTGCAAAACCAAACCACAAGA-3'  Reverse: 5'- CCCAGGTAACCCTTAAAGTCC-3' |
| *MCP1* | NMˍ002982.4 | 58 ºC | Forward: 5'-CTCGCGAGCTATAGAAGAATC-3' Reverse: 5'-TGTGGAGTGAGTGTTCAAGT-3' |
| *TL3* | NMˍ003265.3 | 60 ºC | Forward: 5'-ACTCCACCTCACTATCATGG-3' Reverse: 5'-TCCCAGACCCAATCCTTATC-3' |
| *Β-actin* | NMˍ001101.5 | 63 ºC | Forward: 5'-GTTGCTATCCAGGCTGTGCT-3'  Reverse: 5'-AGGTAGTCAGTGAGGTCCCG-3' |
| *PMSB6* | NMˍ002798.3 | 76 ºC | Forward: 5'-CGGGAAGACCTGATGGCGGGA-3'  Reverse: 5'-TCCCGGAGCCTCCAATGGAAA-3' |
